# Assessing data quality on fetal brain MRI reconstruction: a multi-site and multi-rater study⋆

**DOI:** 10.1101/2024.06.28.601169

**Authors:** Thomas Sanchez, Angeline Mihailov, Yvan Gomez, Gerard Martí Juan, Elisenda Eixarch, András Jakab, Vincent Dunet, Mériam Koob, Guillaume Auzias, Meritxell Bach Cuadra

## Abstract

Quality assessment (QA) has long been considered essential to guarantee the reliability of neuroimaging studies. It is particularly important for fetal brain MRI, where unpredictable fetal motion can lead to substantial artifacts in the acquired images. Multiple images are then combined into a single volume through super-resolution reconstruction (SRR) pipelines, a step that can also introduce additional artifacts. While multiple studies designed automated quality control pipelines, no work evaluated the reproducibility of the manual quality ratings used to train these pipelines. In this work, our objective is twofold. First, we assess the inter- and intra-rater variability of the quality scoring performed by three experts on over 100 SRR images reconstructed using three different SRR pipelines. The raters were asked to assess the quality of images following 8 specific criteria like blurring or tissue contrast, providing a multi-dimensional view on image quality. We show that, using a protocol and training sessions, artifacts like bias field and blur level still have a low agreement (ICC below 0.5), while global quality scores show very high agreement (ICC = 0.9) across raters. We also observe that the SRR methods are influenced differently by factors like gestational age, input data quality and number of stacks used by reconstruction. Finally, our quality scores allow us to unveil systematic weaknesses of the different pipelines, indicating how further development could lead to more robust, well rounded SRR methods. Our rating protocol is made available at https://doi.org/10.5281/zenodo.15696638.

## 1 Introduction

Image quality assessment (QA) is critical to enforce reliability, generalization and reproducibility of neuroimaging studies [1, 2]. Reproducible QA by human raters is particularly critical and challenging in fetal brain MRI examinations. QA is more challenging in fetal MRI than in postnatal acquisitions: exams are typically carried out by acquiring several stacks of 2D fast-spin echo T_2_-weighted (T2w) images with thick slices in orthogonal orientations [3, 4]. While this protocol was designed to minimize the impact of motion during the acquisition, the resulting images can still be severely affected by artifacts like inter-slice motion, signal drops or bias field inhomogeneity [4]. By design, a single fetal MRI acquisition thus corresponds to multiple stacks of 2D images in comparison to a single T1/T2 weighted volume to be assessed for QA in postnatal protocols. QA is critical for fetal MRI because image quality of the original acquisitions affects in a complex manner each of the image processing steps which aim at computing an artifact-free, motion-corrected, 3D volume with isotropic resolution (Figure 1A) from the stacks of 2D images. While these super-resolution reconstruction (SRR) pipelines [5–8] have been designed to compensate for the artifacts affecting the input stacks of 2D images, their processing can converge towards a sub-optimal solution, yielding a SRR 3D volume of insufficient quality for downstream applications, such as biometry measurements or tissue segmentation (Figure 1B). QA for fetal MRI is thus particularly critical when considering potential translation in clinical routine, where the variations in image quality might impact the measurement of interest for diagnostics at the individual level.

**Fig. 1.**
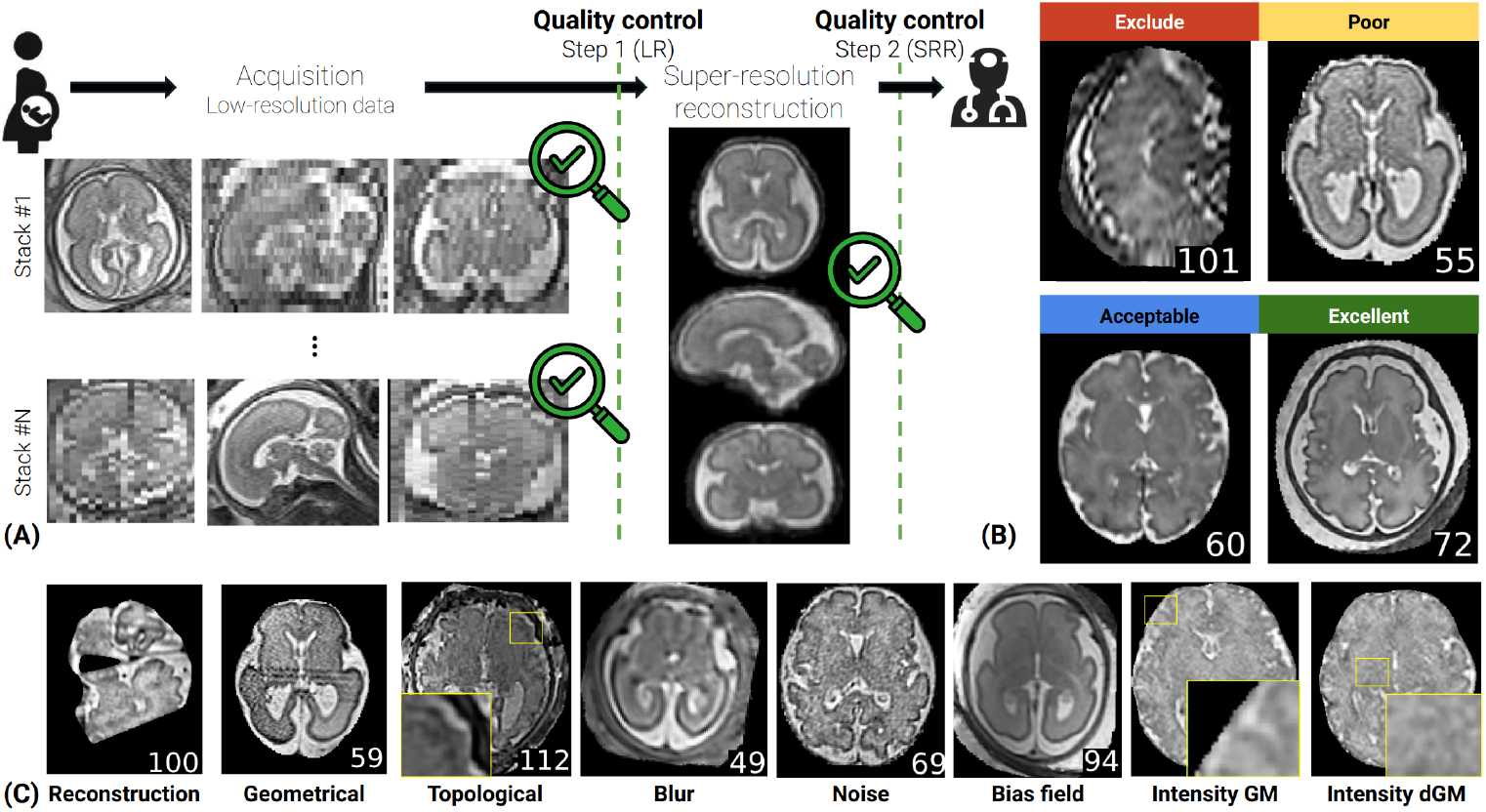
**(A)** Illustration of the data acquisition and reconstruction in fetal brain MRI. Stacks of 2D images are acquired at multiple orientations and combined into a single 3D volume using super-resolution reconstruction techniques. Quality control checks are implemented on the stacks of 2D images (Step 1) and on the SRR volume (Step 2). **(B)** SRR volumes with different quality scores. **(C)** Example of the different artifacts rated in this study.

Some works have thus introduced automated QA methods for either the clinical acquisition of 2D stacks [9–14] or the output SRR volume [15, 16]. Most of them rely on a binary include/exclude criterion, and train a supervised model to identify and exclude poor quality data.

However, most of these works rely on QA scores from a single rater, which runs the risk of developing biased models that overfit to a given rater [17], potentially forfeiting the very objectivity that is sought in automated quality assessment [18]. Multi-rater dataset are commonly developed in segmentation tasks, as they allow to quantify inter-rater variability, providing a bound on the expected performance of a given method, but enable also to build reliable ground truth data [19–21]. Inter-rater QA variability has been estimated on adult MRI [18, 22, 23], showing a large variability across raters, but these works generally attempt to give a global quality score, and rarely focus on rating the degree of importance of specific artifacts. While only a single work reported inter-rater reliability in the context of 2D T2w stacks QA [13], this has never been studied in the context of fetal brain SRR.

In this work, we address the three following open questions: **(1)** How reliable are quality scores on fetal brain SR reconstructions within and across raters? **(2)** What are the artifacts that arise in different SRR algorithms and how can we use these insights to improve the quality of SRR volumes? **(3)** How is the quality of SRR volumes connected to variables like gestational age, number of stacks used in reconstruction and quality of the input LR T2w stacks?

### 2 Methodology

### 2.1 Data

We reconstructed 105 SRR volumes from the 2D clinical series of 21 subjects, acquired across three different scanners and 3 clinical centers, BCNatal (n=8), Lausanne University Hospital (CHUV) (n=11) and University Children’s Hospital Zürich (Kispi) (n=2). The data were acquired on different Siemens (BCNatal and CHUV) and General Electrics (Kispi) scanners at 1.5T (n=14) and 3T (n=7). 15/21 subjects are pathological (5 intrauterine growth restrictions, 3 corpus callosum agenesis (CCA), 3 partial CCA, 3 ventriculomegaly, 1 cytomegalovirus). Details on the input data resolution, reconstruction resolution, gestational age and number of low-resolution stacks available are provided in the supplementary Table S1. The corresponding local ethics committees independently approved the studies under which data were collected, and all participants gave written informed consent. Each subject was then reconstructed using three state-of-the-art reconstruction pipelines: SVRTK [5, 24, 25], NiftyMIC [7] and NeSVoR [8]. It was reconstructed twice with each method: once using all available LR T2w stacks and a second time after filtering out bad quality stacks using quality ratings from FetMRQC [13], an automated QA pipeline for LR T2w stacks. After discarding duplicates (7 subjects where all stacks are included after QC) and assigning a quality score of 0 to reconstructions that failed to produce an output (5 reconstructions from NiftyMIC), this yielded the total of 105 SR volumes.

### 2.2 Manual quality rating

As previous works often showed a substantial level of disagreement on subjective QA tasks [13, 22], we attempted to build a formal QA rating protocol, where the raters were first asked to rate eight specific artifact criteria (Figure 1C) before giving a global subjective quality score (Figure 1B). The categories are the following:

1. **Full reconstruction:** Is the brain fully reconstructed? Are there holes, or large parts of the brain missing?
2. **Geometrical artifacts:** Did the SRR introduce any non-biological, textured artifact like stripes?
3. **Topological artifacts:** Did the SRR introduce any discontinuity in the cortical gray matter (cGM), or are some parts of the white matter (WM) directly touching the cortical cerebrospinal fluid (CSF)?
4. **Blur:** Is there some blur in the image, especially between WM and cGM?
5. **Noise:** Is there a high level of noise in the image, potential preventing us from seeing the deep gray matter (dGM) clearly?
6. **Bias:** Is there a bias field (smooth intensity inhomogeneity) on the image?
7. **Tissue intensity contrast – WM/cGM/cerebrospinal fluid:** Is the contrast sufficient? Do we see well the cGM?
8. **Tissue intensity contrast – WM/dGM:** Is the contrast sufficient? Do we see well the sub-regions in the dGM?

The first artifact is a binary rating, and the rest are continuous, in the form of sliders with values between 0 and 3 or 4, as shown on Figure S1). After rating each of these categories, the raters gave a continuous global quality score, with values corresponding to four categories [0, 1[= exclude, [1, 2[= poor, [2, 3[= acceptable and [3, 4[= excellent. The three raters (R1, R2, R3) underwent a training session on 25 SRR volumes not considered in this work (external to the dataset mentioned above), discussed their results and refined common criteria before rating them again. After this training session, they were asked to manually score the 105 SRR volumes twice, with the cases randomly shuffled. In total, we collected 105 volumes *×* 3 raters *×* 2 repetitions = 630 ratings.

### 2.3 Experiments

#### 1 SRR rating variability

We analyzed the ratings of the 105 SRR volumes by the three raters. We measured inter-rater variability for each the eight artifact ratings as well as the global socre using intraclass correlation coefficient ICC(A,1) (R package irr v.0.84.1) [26]. We did this computation on both repetitions. To measure intra-rater-reliability, we used Lin’s concordance correlation coefficient (CCC) (R package DescTools v.0.99.54) [27].

#### 2 Systematic artifacts of SRR methods

We evaluated the quality of the three SRR algorithms considered by comparing their eight quality scores. We first carried out a Friedman test (the non-parametric equivalent of a repeated measures ANOVA) with two degrees of freedom for each quality score, and corrected the resulting p-values for multiple comparisons using Bonferroni correction. When the p-value was below *α* = 0.05, we ran a pair-wise Wilcoxon signed rank test to identify pairs that showed statistically significant differences. We used the quality ratings averaged across raters and repetitions as the reference quality scores.

#### 3 Influence of gestational age and input 2D stacks quality on the output SRR volume

To study how gestational age (GA) and the input stacks quality influences the SRR volumes, we computed Spearman rank correlation, and visually investigated extreme outliers. As different SRR algorithms could lead to different trends, we evaluated the correlation individually for each SRR method. The LR T2w quality scores were obtained using FetMRQC [13], and a global input quality score was computed by averaging the quality of each stack used in the reconstruction of a given SRR volume. We used the averaged SRR quality ratings across raters and repetitions as the reference SRR quality score.

## 3 Results

### 3.1 Global SR quality ratings are highly reliable

In Table 1, we show the inter- and intra-rater reliability for the 3 raters, by comparing the scores across the two sessions (the SRR volumes were scored twice).

**Table 1.** Inter- and intra-rater reliability on 105 images rated two times by three raters. Inter-rater reliability was computed using intraclass correlation coefficient ICC(A,1) (squared brackets are 95% CI), intra-rater reliability was computed using Lin’s concordance correlation coefficient (CCC) (with standard deviation across raters). Reliability above 0.8 is shown in blue, and below 0.5 in red.

| Rating | Inter-rater reliability |  | Intra-rater reliability |
| --- | --- | --- | --- |
|  | Round 1 | Round 2 |  |
| Full recon. | 0.77 [0.69,0.83] | 0.80 [0.73,0.85] | 0.94±0.01 |
| Geometric | 0.48 [0.33,0.62] | 0.52 [0.39,0.64] | 0.72±0.10 |
| Topological | 0.64 [0.51,0.74] | 0.62 [0.45,0.74] | 0.74±0.07 |
| Blur | 0.46 [0.32,0.59] | 0.47 [0.32,0.60] | 0.57±0.04 |
| Noise | 0.74 [0.66,0.82] | 0.69 [0.55,0.79] | 0.80±0.03 |
| Bias | 0.40 [0.22,0.56] | 0.50 [0.37,0.62] | 0.55±0.27 |
| Intensity GM | 0.67 [0.55,0.77] | 0.66 [0.50,0.77] | 0.85±0.03 |
| Intensity dGM | 0.65 [0.53,0.75] | 0.68 [0.55,0.78] | 0.85±0.03 |
| Global Quality | 0.89 [0.86,0.92] | 0.89 [0.85,0.92] | 0.94±0.02 |

**Table 2.**
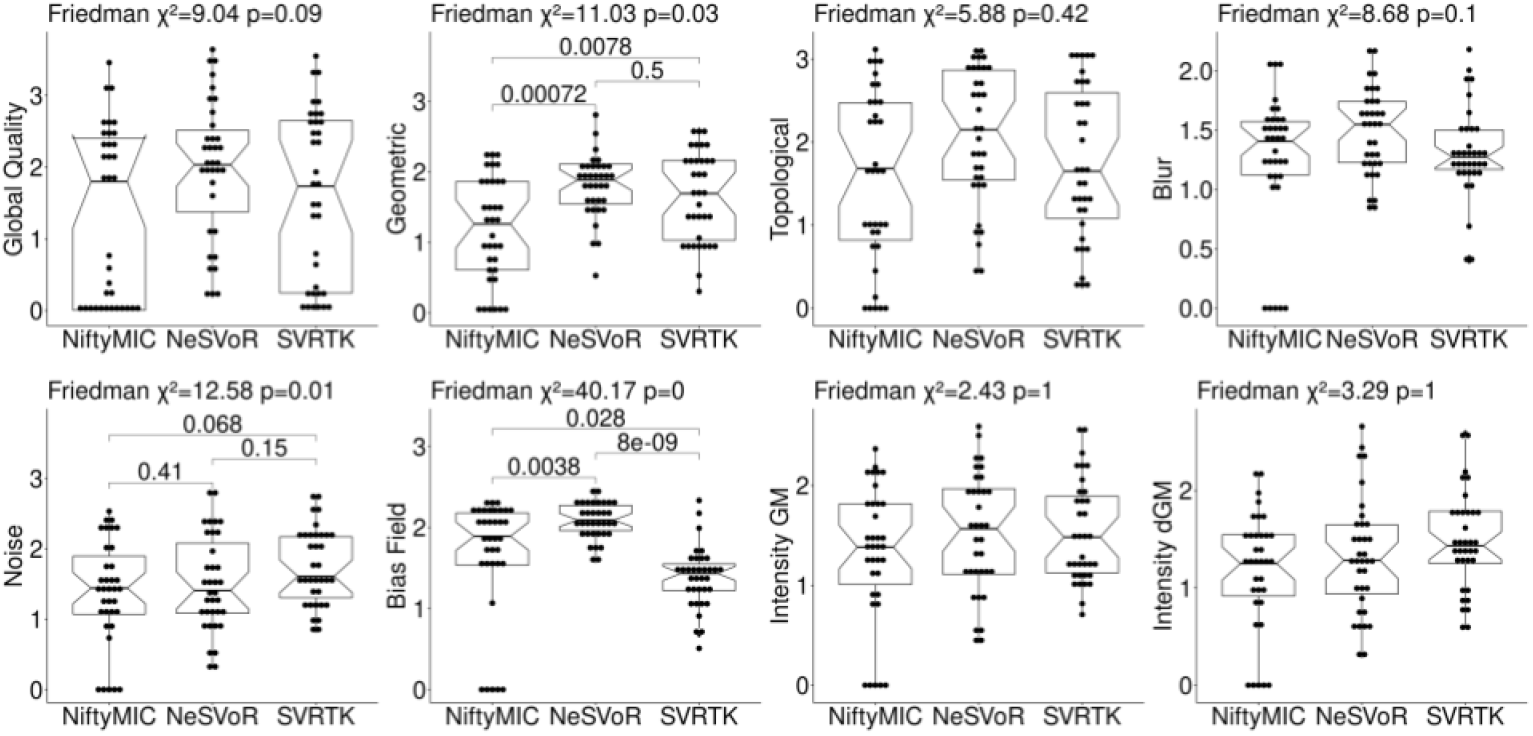
Box plots showing the difference in all quality ratings across the three reconstruction pipelines used in this study.

Although raters underwent a training session, we see that the both inter- and intrarater reliability remain low for several criteria, namely the level of blur in the image (particularly at the CSF/GM/WM interface), and of the level of bias. The inter-rater agreement for the level of geometric artifacts was also low, although the intra-rater reliability was higher, suggesting a systematic discrepancy in scoring across raters. The blur and bias levels correspond to a moderate reliability according to Koo and Li’s interpretation [28]. There is an good-to-excellent level of agreement across raters for the global quality score.

### 3.2 Different SRR methods suffer from different artifacts

Figure 2 shows the box plots for the eight discrete quality ratings, and shows pair-wise Wilcoxon signed rank tests when the Friedman test was found to be significant. A statistical difference was found across the SRR methods for the geometric artifacts, the noise and the bias field rating. For geometric artifacts, NiftyMIC was found to have significantly more artifacts (i.e. lower score) than NeSVoR (*W* = 522, *p* = 0.00072) and SVRTK (*W* = 155, *p* = 0.0078). The pairwise analysis did not return any significant results for the noise rating, likely due to a lack of power of the test. For the bias field, SVRTK was found to have significantly more bias than NiftyMIC (*W* = 450, *p* = 0.028) and NeSVoR (*W* = 615, *p* = 8 *×*10^−9^), and in turn NiftyMIC was found to have significantly more bias than NeSVoR (*W* = 492, *p* = 0.004). This is an expected result as both NiftyMIC and NeSVoR include N4 bias field correction [29] in their pipelines, while SVRTK does not.

**Fig. 2.**
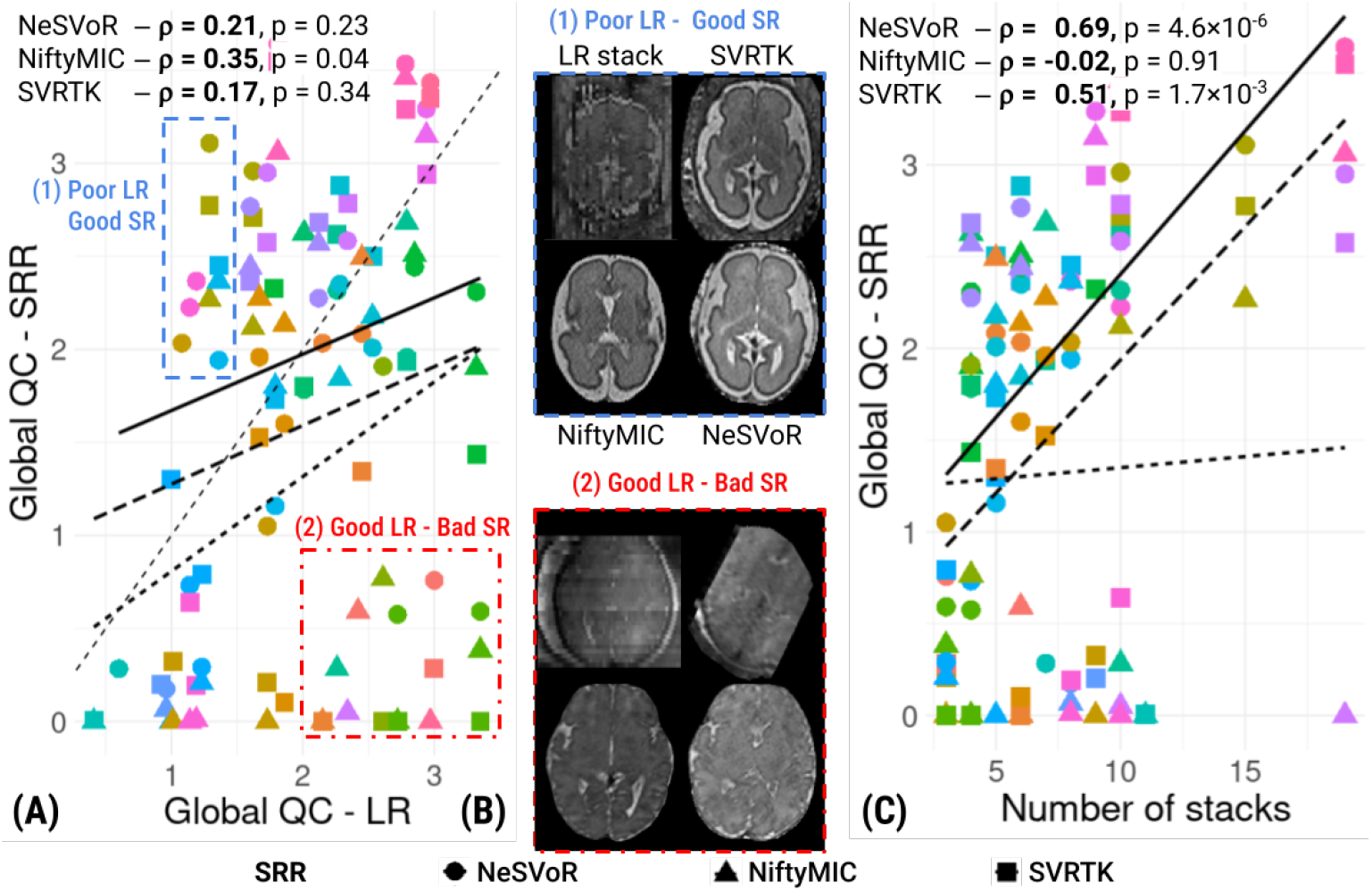
**(A)** Global QA score of the SRR volume as a function of the averaged QA score of the input 2D stacks. Each color shade corresponds to a different subject. The full linear regression line corresponds to NeSVoR, the short dashed one to NiftyMIC and the long dashed line to SVRTK. The Spearman rank correlation coefficient *ρ* is reported for each SRR method. **(B)** Illustration of extreme cases from Figure 2A. *Top box:* example of a case where the LR data were rated of poor quality, but the reconstructions are of very good quality (top left in (A)). *Bottom box:* example of a case where the LR data were rated as high quality, but the reconstructions are of poor quality (bottom right in (A)). **(C)** Global QA score of the SRR volume as a function of the number of stacks provided as input.

### 3.3 SRR methods depend differently on LR T2w quality, gestational age and number of stacks used in reconstruction

Figure 2A shows the correlation between the averaged quality rating of the 2D stacks and the SRR volume quality. The Spearman rank correlations are quite low, with *ρ*_NeSVoR_ = 0.22, *ρ*_NiftyMIC_ = 0.35, *ρ*_SVRTK_ = 0.17. There are two types of outliers: region (1) shows 16 cases, out of 53 cases that have been labeled as having very good quality 2D stacks, where the reconstruction was deemed unusable (quality below 1) – they greatly impact the correlation, as removing them changes correlation to *ρ* = 0.53; region (2) shows eight opposite cases, where the input 2D stacks were deemed of poor quality (16 cases in total ∈ [1, 1.5[), but the SRR was between acceptable and excellent quality. A picture of the 2D stacks and SRR with all three SRR methods is displayed in Figure 2B. We can see that among the cases with good scores for 2D stacks and failed SRR, the score for 2D stacks was excellent for 8/16 stacks because they displayed very little motion, although their contrast was low and bias field was high. These cases lead to very poor reconstruction. The rest of the cases with a good quality for 2D stacks and bad SRR feature one subject with ventriculomegaly, where NiftyMIC and SVRTK failed, two failures by SVRTK and four failures from NiftyMIC. We also observe that these two regions feature SRR obtained with a very different number of stacks. Region (1) has *n*_stacks_ = 10.6 *±* 3.4 while region (2) only has *n*_stacks_ = 5.19 *±* 2.6. This is consistent with Figure 2C, where we observed that NeSVoR and SVRTK benefit a lot from having more stacks available, with a correlation *ρ* = 0.69 and 0.51 respectively, while this is not the case for NiftyMIC (*ρ* = −0.02). We also investigated the effect of GA on SRR volume quality, and detailed results are available on Figure S3 in the Supplementary Material. The Spearman rank correlation shows different trends for the different SRR methods. NeSVoR and SVRTK show a statistically significant negative correlation (*ρ* = −0.38, *ρ* = −0.44) while NiftyMIC shows a small positive, non-significant correlation. This is not surprising for NeSVoR and SVRTK, as subjects at older GAs have a more developed brain with more subtle structures, and as a result, similar artifacts can be perceived as impacting more the quality compared to younger GAs. The opposite trend in NiftyMIC is caused by many subjects where NiftyMIC’s outlier rejection excluded most of the available data, leading to a SRR of unacceptable quality.

## 4 Discussion and conclusion

In this study, we quantified for the first time the intra- and inter-rater variability in SRR quality assessment. We then used these ratings to better understand the differences between NeSVoR, NiftyMIC and SVRTK in terms of their susceptibility to the artifacts considered. Finally, we studied how external factors like the quality of the input data, the number of stacks used for reconstruction and the gestational age influenced the behavior of these SRR pipelines. **Increasing further the reliability of QA scores**. Our results show that excellent reliability can be achieved for global quality scoring across three raters, with good reliability on specific criteria relating to the contrast across tissues and noise levels. However we also identified blurriness and bias field as difficult to assess by raters, with the lowest inter- and intra-rater reliability in the study. This is in line with previous work [13] that found almost no correlation between raters on rating of the level of bias field in LR images. Additional visualization tools based on a difference map between the original image and a de-biased image could potentially help human raters in this task. Such tools have been implemented for adult brain MRI QC, where visual reports feature enhancement of the noise around the head that helps to identify various motion-related artifacts [22].

### Improving quality control in low-resolution data

As we have seen in Experiment 3, there are many outliers that have been assigned a good LR quality, but lead to an unusable SRR. In particular, input LR data with low contrast were rated very highly, while they systematically led to a failed reconstruction. This should be taken into account in future works on LR quality rating: while in LR quality rating, one easily notices and penalizes artifact due to motion (signal drops or staircase-like motion through plane), they can be addressed more reliably than a very low contrast input.

### Improving SRR reconstruction methods

Our results from Experiments 2 and 3 show potential avenues for improving the performance of SRR pipelines. In particular, Figure 2 shows that SVRTK leads to volumes with a strong bias field. Running a bias field correction algorithm like N4 [29] could then be considered to address this issue. NiftyMIC was shown to exhibit significant geometric artifacts, caused by its outlier rejection module excluding too many slices. This results in several cases where the majority of data is discarded, preventing it from benefiting from using more stacks, as opposed to SVRTK and NeSVoR. Tuning the exclusion parameter more carefully could be an avenue to improve NiftyMIC’s performance. In general, simulated data could be of help to tune these parameters in a quantitative way, complementing manual annotations [30].

### Future perspectives

Future extensions of this work will focus on further improving the quality and robustness of SRR methods. In particular, we will investigate how we can use the quality assessment at the input level to maximize the SRR quality. From the clinical perspective, automating fetal brain MRI QA and integrating it in the clinical workflow would reduce the burden of manually processing acquired images and would also lead to more automated MRI-to-PACS pipelines for fetal MRI analysis.

## Supplementary material

**Fig. S1.**
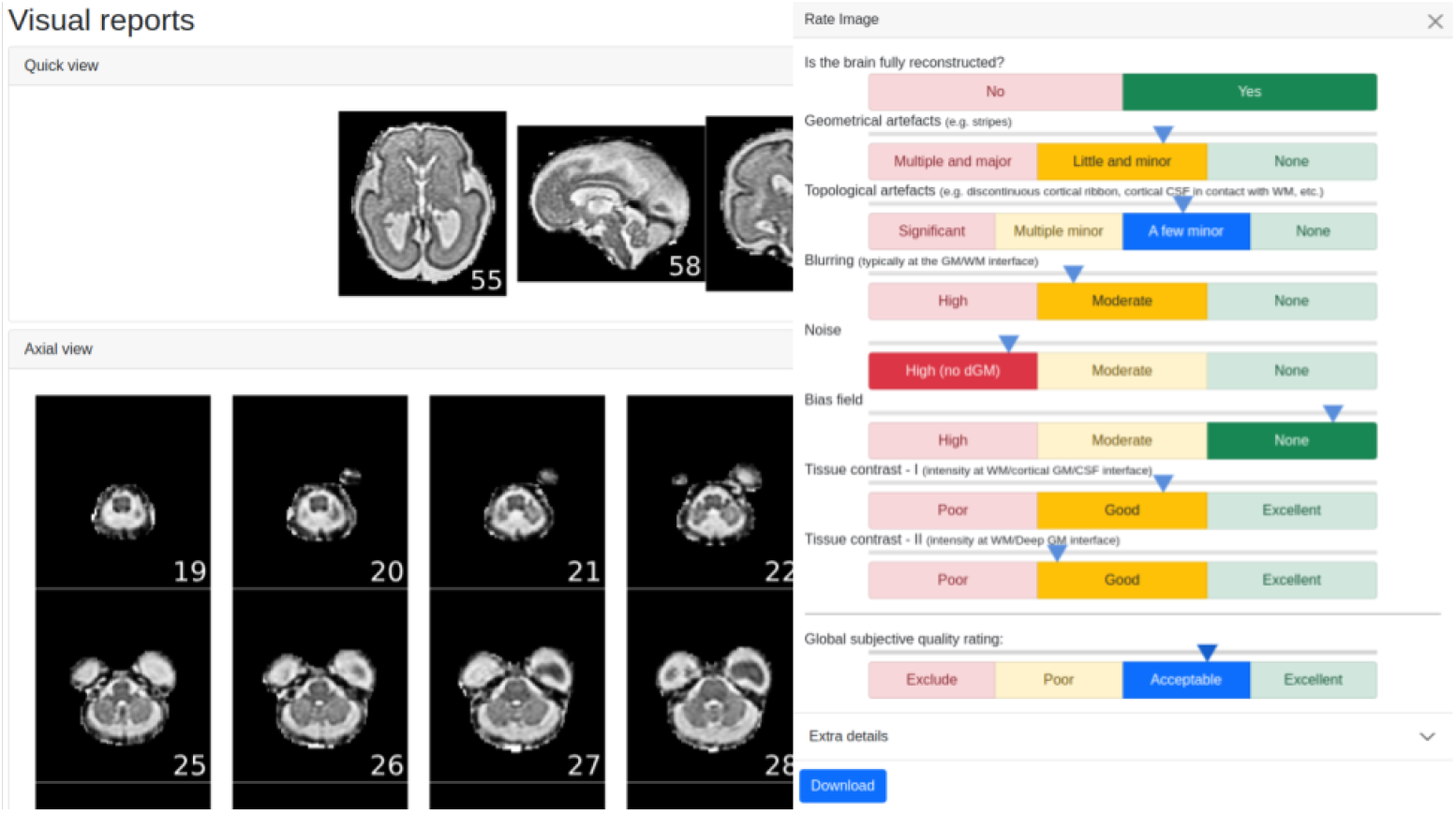
Example of a manual QC report used in the study. The report display a quick view with a slice in axial, sagittal and coronal orientations, and then shows all the slices from all orientations. The rating panel contains categories of artifact severity for each entry. Geometrical artifacts, blurring, noise, bias field, tissue contrasts I (WM/cGM/CSF) and II (WM/dGM) are rated on a scale from 0 (very severe artifact/poor quality) to 3 (no artifact/excellent quality). Topological artifacts and global quality score are rated in a scale from 0 to 4.

**Table S1.**
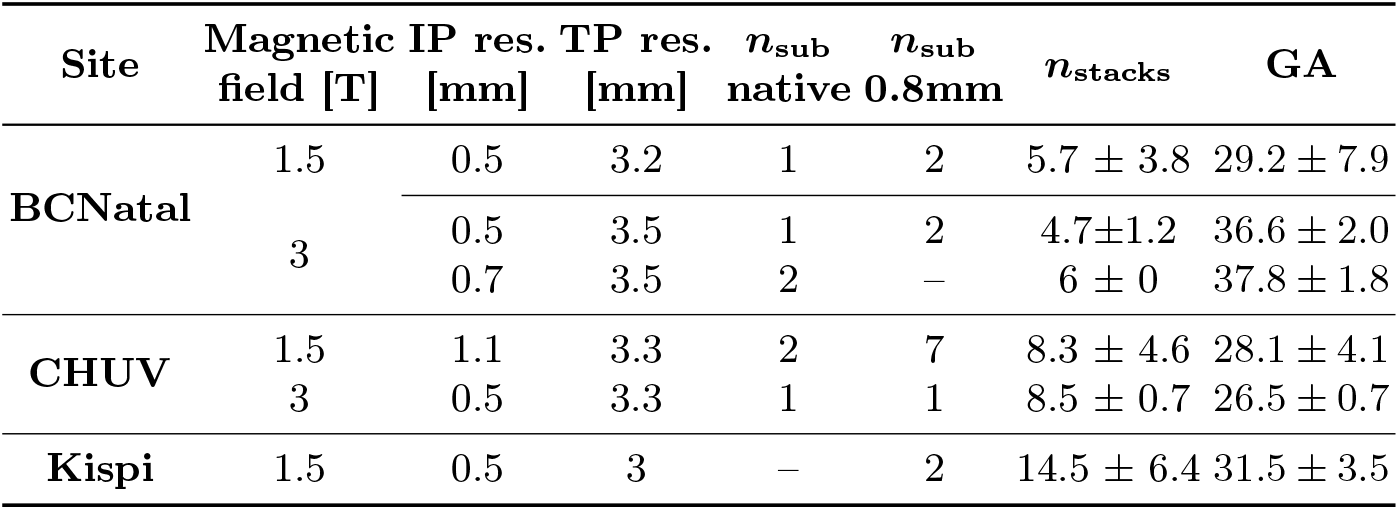
Data description. *n*_sub_ native and *n*_sub_ 0.8mm refer to the number of subjects reconstructed either at their native resolution (0.5mm or 1.1mm isotropic) or at 0.8mm isotropic. This allows to account for the bias that different target resolutions might introduce in the data. The BCNatal data contain 5 subjects with intrauterine growth restriction (IUGR) and 3 with corpus callosum agenesis (CCA) pathological subjects and the CHUV data contain 3 subjects with ventriculomegaly, 3 with (partial) CCA, 1 with cytomegalovirus (CMV) and 4 controls.

**Fig. S2.**
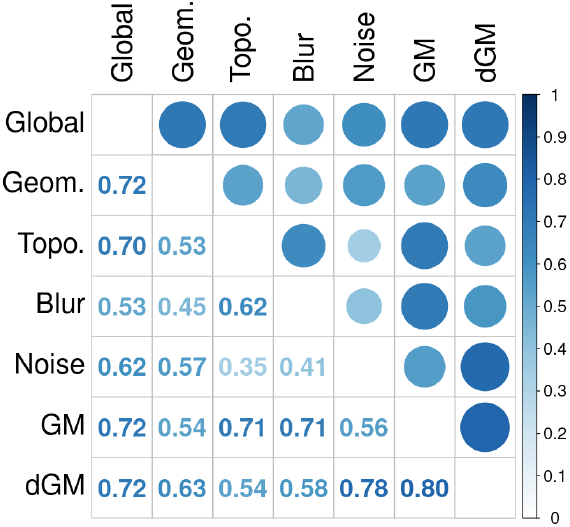
Correlation matrix between the different rating criteria across all raters.

**Fig. S3.**
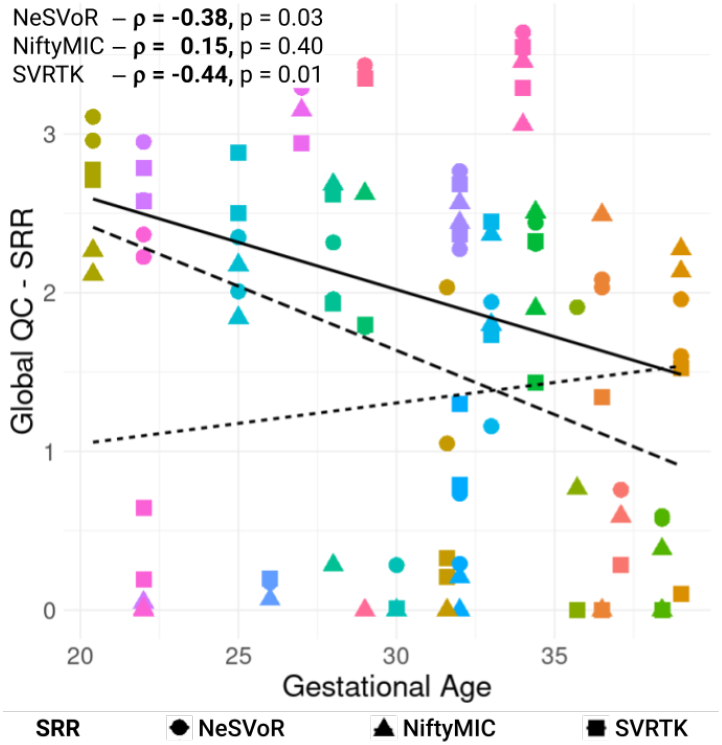
Global QA score of the SRR volume as a function of the gestational age. Each color shade corresponds to a different subject. The full linear regression line corresponds to NeSVoR, the short dashed one to NiftyMIC and the long dashed line to SVRTK. The Spearman rank correlation coefficient *ρ* is reported for each SRR method.

## Notes

⋆ TS, GA, AM, GMJ were supported by the Era-Net Neuron MULTIFACT project, under local project numbers SNSF 31NE30_203977 (TS), ANR-21-NEU2-0005 (GA,AM), PCI2021-122044-2A (GMJ). GA, AM acknowledge support from the SulcalGRIDS project (ANR-19-CE45-0014). YG acknowledges support from the SICPA foundation and EE is supported by the Instituto de Salud Carlos III (ISCIII)(AC21_2/00016). AJ is supported by the Prof. Max Cloetta Foundation, EMDO Foundation and Vontobel Foundation. TS and MBC acknowledge access to the facilities and expertise of the CIBM Center for Biomedical Imaging, a Swiss research center of excellence founded and supported by CHUV, UNIL, EPFL, UNIGE and HUG.

https://doi.org/10.5281/zenodo.15696638

### Competing Interest Statement

The authors have declared no competing interest.

### Summary of Updates

Add a link to the protocol used for the manual quality rating.

## References

1. A. F. Rosen, D. R. Roalf, K. Ruparel, J. Blake, K. Seelaus, L. P. Villa, R. Ciric, P. A. Cook, C. Davatzikos, M. A. Elliott et al., “Quantitative assessment of structural image quality,” Neuroimage, vol. 169, pp. 407–418, 2018.

2. G. Niso, R. Botvinik-Nezer, S. Appelhoff, A. De La Vega, O. Esteban, J. A. Etzel, K. Finc, M. Ganz, R. Gau, Y. O. Halchenko et al., “Open and reproducible neuroimaging: from study inception to publication,” NeuroImage, p. 119623, 2022.

3. P. Tortori-Donati, A. Rossi, N. Girard, and T. A. Huisman, “Fetal magnetic resonance imaging of the central nervous system,” Pediatric Neuroradiology: Brain, pp. 1219–1253, 2005.

4. A. Gholipour et al., “Fetal MRI: a technical update with educational aspirations,” Concepts in Magnetic Resonance Part A, vol. 43, no. 6, pp. 237–266, 2014.

5. M. Kuklisova-Murgasova et al., “Reconstruction of fetal brain MRI with intensity matching and complete outlier removal,” Medical image analysis, vol. 16, no. 8, pp. 1550–1564, 2012.

6. S. Tourbier, X. Bresson, P. Hagmann, J.-P. Thiran, R. Meuli, and M. B. Cuadra, “An efficient total variation algorithm for super-resolution in fetal brain mri with adaptive regularization,” NeuroImage, vol. 118, pp. 584–597, 2015.

7. M. Ebner et al., “An automated framework for localization, segmentation and super-resolution reconstruction of fetal brain MRI,” NeuroImage, vol. 206, p. 116324, 2020.

8. J. Xu, D. Moyer, B. Gagoski, J. E. Iglesias, P. E. Grant, P. Golland, and E. Adalsteinsson, “NeSVoR: Implicit neural representation for slice-to-volume reconstruction in MRI,” IEEE Transactions on Medical Imaging, 2023.

9. S. Lala et al., “A deep learning approach for image quality assessment of fetal brain MRI,” in Proceedings of the 27th Annual Meeting of ISMRM. Montréal, Québec, Canada, 2019, p. 839.

10. J. Xu, S. Lala, B. Gagoski, E. Abaci Turk, P. E. Grant, P. Golland, and E. Adalsteinsson, “Semi-supervised learning for fetal brain mri quality assessment with roi consistency,” in Medical Image Computing and Computer Assisted Intervention–MICCAI 2020: 23rd International Conference, Lima, Peru, October 4–8, 2020, Proceedings, Part VI 23. Springer, 2020, pp. 386–395.

11. L. Liao, X. Zhang, F. Zhao, T. Zhong, Y. Pei, X. Xu, L. Wang, H. Zhang, D. Shen, and G. Li, “Joint image quality assessment and brain extraction of fetal MRI using deep learning,” in Medical Image Computing and Computer Assisted Intervention–MICCAI 2020: 23rd International Conference, Lima, Peru, October 4–8, 2020, Proceedings, Part VI 23. Springer, 2020, pp. 415–424.

12. T. Sanchez, O. Esteban, Y. Gomez, E. Eixarch, and M. Bach Cuadra, “FetMRQC: Automated quality control for fetal brain MRI,” in Perinatal, Preterm and Paediatric Image Analysis. Cham: Springer Nature Switzerland, 2023, pp. 3–16.

13. T. Sanchez et al., “FetMRQC: A robust quality control system for multi-centric fetal brain MRI,” Medical Image Analysis, p. 103282, 2024.

14. W. Zhang, X. Zhang, L. Li, L. Liao, F. Zhao, T. Zhong, Y. Pei, X. Xu, C. Yang, H. Zhang et al., “A joint brain extraction and image quality assessment framework for fetal brain MRI slices,” NeuroImage, p. 120560, 2024.

15. A. Largent et al., “Image quality assessment of fetal brain MRI using multi-instance deep learning methods,” Journal of Magnetic Resonance Imaging, vol. 54, no. 3, pp. 818–829, 2021.

16. N. Rubert, D. M. Bardo, J. Vaughn, P. Cornejo, and L. F. Goncalves, “Data quality assessment for super-resolution fetal brain MR imaging: A retrospective 1.5 T study,” Journal of Magnetic Resonance Imaging, vol. 54, no. 4, 2021.

17. O. Shwartzman et al., “The worrisome impact of an inter-rater bias on neural network training,” arXiv preprint 1906.11872, 2019.

18. J. Monereo-Sánchez et al., “Quality control strategies for brain MRI segmentation and parcellation: Practical approaches and recommendations-insights from the Maastricht study,” Neuroimage, vol. 237, p. 118174, 2021.

19. Ž. Lesjak et al., “A novel public MR image dataset of multiple sclerosis patients with lesion segmentations based on multi-rater consensus,” Neuroinformatics, vol. 16, pp. 51–63, 2018.

20. A. S. Becker, K. Chaitanya, K. Schawkat, U. J. Muehlematter, A. M. Hötker, E. Konukoglu, and O. F. Donati, “Variability of manual segmentation of the prostate in axial T2-weighted MRI: a multi-reader study,” European journal of radiology, vol. 121, p. 108716, 2019.

21. L. Joskowicz, D. Cohen, N. Caplan, and J. Sosna, “Inter-observer variability of manual contour delineation of structures in CT,” European radiology, vol. 29, pp. 1391–1399, 2019.

22. O. Esteban, D. Birman, M. Schaer, O. O. Koyejo, R. A. Poldrack, and K. J. Gorgolewski, “MRIQC: Advancing the automatic prediction of image quality in MRI from unseen sites,” PloS one, vol. 12, no. 9, p. e0184661, 2017.

23. E. T. Klapwijk, F. Van De Kamp, M. Van Der Meulen, S. Peters, and L. M. Wierenga, “Qoala-T: A supervised-learning tool for quality control of FreeSurfer segmented MRI data,” Neuroimage, vol. 189, pp. 116–129, 2019.

24. A. U. Uus, M. Hall, K. Payette, J. V. Hajnal, M. Deprez, M. A. Rutherford, J. Hutter, and L. Story, “Combined quantitative T2* map and structural T2-weighted tissue-specific analysis for fetal brain MRI: Pilot automated pipeline,” in International Workshop on Preterm, Perinatal and Paediatric Image Analysis. Springer, 2023, pp. 28–38.

25. A. U. Uus et al., “Scanner-based real-time 3D brain+ body slice-to-volume reconstruction for T2-weighted 0.55 T low field fetal MRI,” medRxiv, 2024.

26. K. O. McGraw and S. P. Wong, “Forming inferences about some intraclass correlation coefficients.” Psychological methods, vol. 1, no. 1, p. 30, 1996.

27. L. I.-K. Lin, “A concordance correlation coefficient to evaluate reproducibility,” Biometrics, pp. 255–268, 1989.

28. T. K. Koo and M. Y. Li, “A guideline of selecting and reporting intraclass correlation coefficients for reliability research,” Journal of chiropractic medicine, vol. 15, no. 2, pp. 155–163, 2016.

29. N. Tustison, B. Avants, P. Cook, Y. Zheng, A. Egan, P. Yushkevich, and J. Gee, “N4itk: Improved N3 bias correction,” IEEE Transactions on Medical Imaging, vol. 29, no. 6, 2010.

30. P. de Dumast, T. Sanchez, H. Lajous, and M. Bach Cuadra, “Simulation-based parameter optimization for fetal brain MRI super-resolution reconstruction,” in International Conference on Medical Image Computing and Computer-Assisted Intervention. Springer, 2023, pp. 336–346.

